# The Temporal Constraints of the Cerebellum’s Timekeeping

**DOI:** 10.64898/2026.04.10.717630

**Authors:** Kelly Hoogervorst, Lau Møller Andersen

**Author notes:** Corresponding author information: Kelly Hoogervorst.

## Abstract

The cerebellum plays a central role in generating temporal predictions from past sensory regularities, yet the temporal boundaries of this predictive capacity remain unclear. Using magnetoencephalography (MEG), we investigated somatosensory and cerebellar responses to omissions within rhythmic somatosensory stimulation trains across six inter-stimulus intervals (ISIs) ranging from 0.5 to 5.5 seconds. We hypothesised that cerebellar prediction signals would follow a logistic decay pattern, remaining robust at short ISIs before declining beyond a 2–4-second temporal threshold. As a first step, we validated the omission paradigm by confirming the expected SI and SII response pattern to stimulations and the preservation of the SII response to omissions. Cerebellar source reconstruction revealed consistent beta band (14-30 Hz) responses to omissions peaking at 40–50 ms post-omission in right lobule VI, replicating previous findings. Critically, cerebellar activation to omissions was compatible with a logistic decay pattern with increasing ISI, with the inflexion point estimated within the hypothesised 2–4-second window, though precise localisation of this threshold warrants further investigation. Together, these findings establish empirical boundaries for cerebellar temporal prediction, suggesting that the cerebellum operates as a precise but duration-limited internal clock with implications for understanding the brain’s timing mechanisms and their functional consequences for perception.

## 1. Introduction

It is increasingly clear that perception and prediction cannot be fully disentangled, as prediction of future events depends on extracting regularities from past perceptual experience. The cerebellum has been implicated in building so-called forward models that contain information about what, where and when, based on past regularities (Andersen & Dalal, 2024; Ernst et al., 2019; Kilteni et al., 2018). In the temporal domain, the cerebellum has been proposed to also directly perform the relevant timing predictions (Ivry et al., 2002). A remaining question concerns the maximum duration over which the cerebellum can integrate information to build such predictions.

Using magnetoencephalography (MEG), Tesche & Karhu (2000) found early evidence that trains of stimulation elicit cerebellar oscillations up to 2-4 seconds after stimulation. This 2-4-second window of cerebellar oscillatory activity may constrain the timing of action, potentially explaining why humans cannot maintain rhythmic tapping with intervals exceeding 2-3 seconds (Mates et al., 1994; Miyake et al., 2004). By presenting omissions of otherwise expected stimuli within temporally regular trains of somatosensory stimulation, we have previously found evoked responses of the secondary somatosensory cortex (SII) to be elicited by inter-stimuli intervals (ISIs) shorter than or equal to 3 seconds (Andersen & Dalal, 2021; Andersen & Lundqvist, 2019). Furthermore, we have found oscillatory responses in the cerebellar beta band (14-30 Hz) for the omissions, peaking ∼45 milliseconds after the expected, but omitted, stimulus in cerebellar lobule VI and cerebellar crus I for an ISI of 1.5 seconds (Andersen & Dalal, 2021, 2024). Even though cerebellar source reconstruction in MEG is seemingly challenged by the cerebellum’s distance from sensors, complex folding, and cancellation effects from opposing cortical surfaces, Samuelsson et al. (2020) have conclusively shown that MEG is sufficiently sensitive to reliably detect cerebellar activations. In the current study, we use MEG to investigate the degree to which cerebellar responses depend on the temporal distance between preceding stimuli, i.e. past events. If there is such a dependency, it may be interpretable as reflective of the prediction precision in this region.

We hypothesise that the relationship between ISI and the cerebellar beta-band response is of a logistic nature. That is, we expect a baseline range of ISIs that can be timed by the cerebellum, as indicated by a cerebellar beta-band response to omissions. This baseline range should be followed by a drop in the magnitude of the beta-band response to omissions for ISIs longer than 2-4 seconds (Tesche & Karhu, 2000), because the ISI can no longer be timed precisely by the cerebellum.

Using a generalised logistic function, we can model this relation using the following equation,

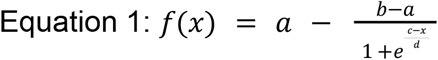

where *x* is the ISI in seconds, *a* is the (expected) upper asymptote, *b* is the (expected) lower asymptote, *c* is the inflexion point, i.e. where the acceleration, the second derivative, changes its sign, and *d* is an indicator of the steepness of the curve; as *d* approaches 0, *f*(*x*) goes towards a step function, and as *d* goes towards infinity, *f*(*x*) goes towards a linear function. According to the hypothesis, the inflexion point, *c*, should be around 2-4 s (Tesche & Karhu, 2000).

Before examining cerebellar responses, we first validate the omission paradigm by confirming that tactile stimulations elicit the canonical SI (∼50 ms) and SII (∼135 ms) responses, and that omissions preserve the SII response while the early SI component is absent, replicating (Andersen & Lundqvist, 2019). Regarding the cerebellum, we hypothesise the neural beta-band response to omissions is close to the estimated *a* value for ISIs < 2 seconds, whereas it should be close to the estimated *b* value for ISIs > 4 seconds. The inflexion point, *c*, is hypothesised to lie between 2-4 seconds (Tesche & Karhu, 2000).

## 2. Methods

### 2.1 Participants

Data from 26 healthy, right-handed participants were included in the analysis (14 F/12 M, age = 24.9 years, σ = 3.2). Participants were recruited through advertisements on social media and a local database for research participation. Before participating, participants provided their informed consent. An initial sample of 32 participants took part in the experiment. One participant was excluded from the analysis because they did not complete the study. The data of eight participants were subject to a technical error that rendered their data unusable; of these, three returned for an additional session to provide usable data. Participants were remunerated for their participation following standard rates in Denmark. The study was approved by the Institutional Review Board at the Danish Neuroscience Center at Aarhus University.

### 2.2 Stimuli and Procedure

Participants received tactile stimuli in the form of an electric current that was generated by two ring electrodes powered by an electric current generator (Stimulus Current Generator, DeMeTec GmbH, Germany). Current strength was determined individually by self-reported measures of detectability, resulting in an average current strength of 4.37 mA (SD = 1.07 mA). Stimuli were presented in trains consisting of electrical pulses, with each pulse lasting 100 µs. The ISI within each train was set to one of six values: 0.497, 1.397, 2.597, 3.397, 4.597, or 5.397 seconds. These intervals were chosen to prevent ISIs from being multiples of one another. Each stimulus train consisted of either three or four pulses, followed by an omission at the fourth or fifth position, respectively (see Figure 1). This unpredictability was introduced to prevent expectation effects. Omission positions were not distinguished in subsequent analyses. For each ISI, 50 trains were presented (300 total), delivered in blocks of 10 consecutive repetitions before shifting to a new ISI. Omissions were equally distributed between the two potential positions (4th or 5th stimulus) and pseudorandomised across blocks to reduce predictability. While participants received the stimuli, they were shown a nature documentary with sound to keep their attention off the applied stimuli.

**Figure 1.**
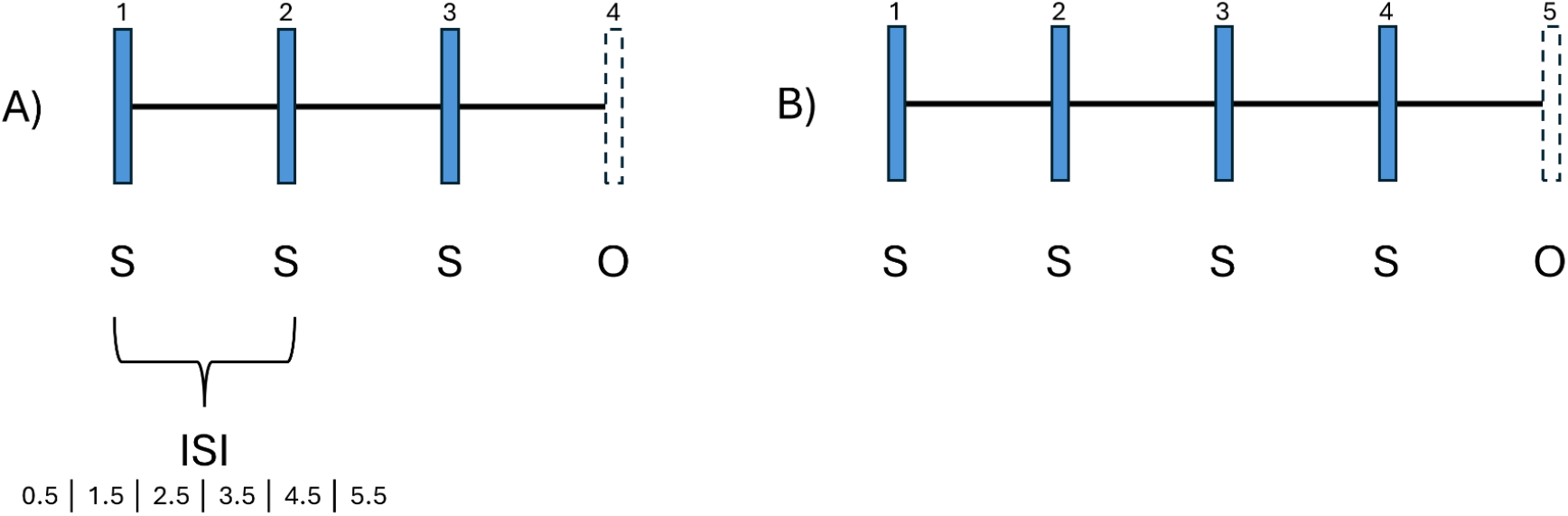
Visualisation of the task paradigm. A) Representation of the first kind of stimulus train used in the experiment, with an omission after three stimuli. At the bottom of the figure, the different ISIs used in the paradigm are denoted in seconds. B) Representation of the second kind of stimulus train used in the experiment, with an omission after four stimuli. Both sequences were represented equally in the paradigm.

### 2.3 Preparation of participants

Prior to scanning, each participant had 11 electrodes placed. This included a pair of electrooculography (EOG) electrodes above and below the right eye, another EOG pair on the temples, a pair of electrocardiogram (ECG) electrodes on both collar bones, a pair of electromyography (EMG) electrodes on each splenius muscle, and a ground electrode on the inside of the right wrist. Additionally, four head-position indicator (HPI) coils were placed: two behind the ears and two on the forehead. Subsequently, each participant’s head shape was digitised using a Polhemus FASTRAK. This was done by first digitising three fiducial points: the nasion and the left and right pre-auricular points, followed by the positions of the HPI-coils. Furthermore, the head shape of each participant was mapped by placing digital points on the head and face. Inside the scanner room, participants were further equipped with a respiration belt and ring electrodes around the top knuckle of the index finger. The first electrode was placed just below the fingernail, and the second approximately 10 mm below that. Participants were then placed in the MEG in supine position, and pillows were placed to ensure comfort and to prevent neck tension. Before the start of the experiment, the stimulus current was individually calibrated. Participants were instructed that the stimulation should be clearly detectable, but without irritation or pain. The baseline current was 2 mA, and increased by 0.5 until the participant reported it as irritating. Then, the stimulation was decreased in increments of 0.5 until the participant felt comfortable with the stimulation. Finally, a train of 10 stimulations was given to simulate the experimental design and confirm the stimulus strength for the experiment.

### 2.4 Acquisition of data

All data were collected at Aarhus University Hospital (AUH), Aarhus, Denmark. MEG data were collected using an Elekta Neuromag TRIUX system located in a magnetically shielded room (Vacuumschmelze Ak3b) with a sampling rate of 1,000 Hz. Online filtering was applied during acquisition, including a low-pass filter at 330 Hz and a high-pass filter at 0.1 Hz. High-resolution sagittal T1-weighted 3D structural images were obtained for each participant during a separate session using a Siemens Magnetom Prisma 3T MRI scanner. Imaging parameters included: 1 mm isotropic voxel size; field of view of 256 × 256 mm; 192 slices with a slice thickness of 1 mm; bandwidth of 290 Hz/pixel; flip angle of 9°; inversion time (TI) of 1,100 ms; echo time (TE) of 2.61 ms; and repetition time (TR) of 2,300 ms.

### 2.5 MEG processing

All MEG processing and analysis was performed in MNE-Python (Gramfort et al., 2013), following a procedure similar to that described in (Andersen, 2018). Before data processing, power spectral densities were manually inspected to identify bad sensors to be excluded from analysis. This resulted in an average of 2 (σ = 0.65) excluded channels per participant.

For analysis of evoked responses, the data was low-pass filtered at 40 Hz (one-pass, noncausal, finite impulse response; zero phase; upper transition bandwidth: 10.00 Hz (−6 dB cutoff frequency: 45.00 Hz; filter length 331 samples; a 0.0194 passband ripple and 53 dB stopband attenuation)), epoched from 100 ms pre-stimulus or -omission to 350 ms post-stimulus or -omission, and baseline-corrected (-100 ms – 0 ms). For each ISI condition, epochs were averaged for each participant for stimulations and omissions separately. Finally, averages were computed as a grand average across all participants.

For analysis in the time frequency domain, the data were band-pass filtered to include the beta band (14–30 Hz; lower transition bandwidth: 3.50 Hz (−6 dB cut-off frequency: 12.25 Hz); upper transition bandwidth: 7.50 Hz (−6 dB cut-off frequency: 33.75 Hz); filter length 943 samples; passband ripple: 0.0194; stopband attenuation: 53 dB)). Subsequently, a Hilbert transform was performed, the data were epoched between 250 ms pre- and post-stimulus or pre- and post-omission, and demeaned using the whole window (-250 ms to 250 ms).

### 2.6 Source reconstruction

First, brain segmentation was performed on T1 images using FreeSurfer (Fischl, 2012) to generate individualised boundary element method (BEM) models. Co-registration between structural MRI and MEG data was achieved by aligning fiducial points (nasion, left/right preauricular) identified on T1 scans with corresponding Polhemus-digitised head points acquired during MEG sessions. A volumetric source space with 7.5 mm grid spacing (∼4,000 sources/subject) was defined using FreeSurfer’s watershed algorithm. Forward models were computed by integrating co-registered data, source space, and single-layer BEM surfaces.

Neural activity was reconstructed using a linearly constrained minimum variance (LCMV) beamformer with unit-noise-gain normalisation, processed separately for evoked and time frequency analyses (Sekihara et al., 2008). Data covariance matrices were estimated from 0 ms to 350 ms post-stimulus for the evoked responses. For beta band responses, data covariance matrices were estimated for the full epoch without regularisation. Only magnetometer data was used in the analysis.

### 2.7 Analysis

To verify somatosensory responses to the tactile stimulation, we examined evoked responses averaged across all ISI conditions and stimulus positions. We identified characteristic SI and SII responses by their topographical distribution and latency. For omissions, we assessed whether previously reported SII responses were present in our paradigm. The *Parietal Opercular Cortex* from the *Harvard-Oxford atlas* (Jenkinson et al., 2012) was chosen to reflect SII.

We used the LCMV beamformer to localise cerebellar responses to tactile stimulation and omissions. To replicate previous findings, we examined whether the shortest ISI (0.497 s) elicited significant cerebellar activation in the beta band (14-30 Hz) in cerebellar lobule VI. To test whether cerebellar engagement varies with temporal predictability, we contrasted activation between the shortest (0.497 s) and all ISIs collapsed. We expected this ISI, 0.497 s, to have the highest precision of temporal prediction, and thus the highest cerebellar beta-band response.

A cluster permutation test was applied to the peak difference (Maris & Oostenveld, 2007) using 1024 permutations with an initial threshold of *α* = 0.05. To examine whether cerebellar activation followed a step-wise decay pattern across ISIs, we extracted mean activation from the identified cerebellar peak for each condition and participant during the post-omission window. The specific time window was chosen based on the window of the duration of the cluster, if any, informing the rejection of the null hypothesis for the cluster permutation test. *The automated anatomical labelling* atlas was used for the cerebellum (Desikan et al., 2006).

### 2.8 Estimation of the generalised logistic function

We sampled the most likely values for the parameters of the generalised logistic function (Equation 1) using the No-U-Turn Sample (NUTS) algorithm (Homan & Gelman, 2014) with 4 chains, 1,000 tuning steps and 1,000 draws per chain. The parameters, *a, b* and *c*, were sampled on both a subject level and a group level. *d* was held fixed. Furthermore, noise on the observations was sampled as well as a *v* parameter, as the likelihood of the data was modelled as a combination of a normal distribution and an exponential distribution. The priors on *a* and *b* were a normal distribution with µ = 0, σ = 1. The prior on *c* was a log-normal distribution (restricting it to positive values), with µ = 1.10, σ = 0.125 (corresponding to a mean of 3 s). As *c* and *d* could not be simultaneously estimated due to insufficient data around the inflexion point, *d* was fixed at 0.75 to reflect a step-like shift in processing capacity. This allowed us to keep the model focused on the key question of where the cerebellar timing limit falls, i.e. the value of the *c* parameter.

## 3. Results

### 3.1 SI and SII

Sensor space analysis confirmed the validity of the omission paradigm through the somatosensory response patterns for tactile stimulations and omissions (Figure 2). When tactile stimuli were delivered (all conditions and individual stimulations collapsed), the expected sequence of somatosensory responses was observed, with contralateral primary somatosensory cortex (SI) activation peaking at approximately 50 ms post-stimulus, followed by secondary somatosensory cortex (SII) activation at 135 ms (Figure 2A). For omissions, the SI response at 50 ms was absent, while a distinct SII response at 135 ms was preserved (Figure 2B), replicating prior work (Andersen & Lundqvist, 2019). Source reconstruction of the omission response at 135 ms for the longest ISI, which yielded the strongest omission response (Supplementary Figure 1), localised the maximally responding source to the right secondary somatosensory cortex (Figure 2C). The time course of this source confirmed a significant response peaking at 135 ms, exceeding the t-threshold for significance (α = 0.05; Figure 2D).

**Figure 2.**
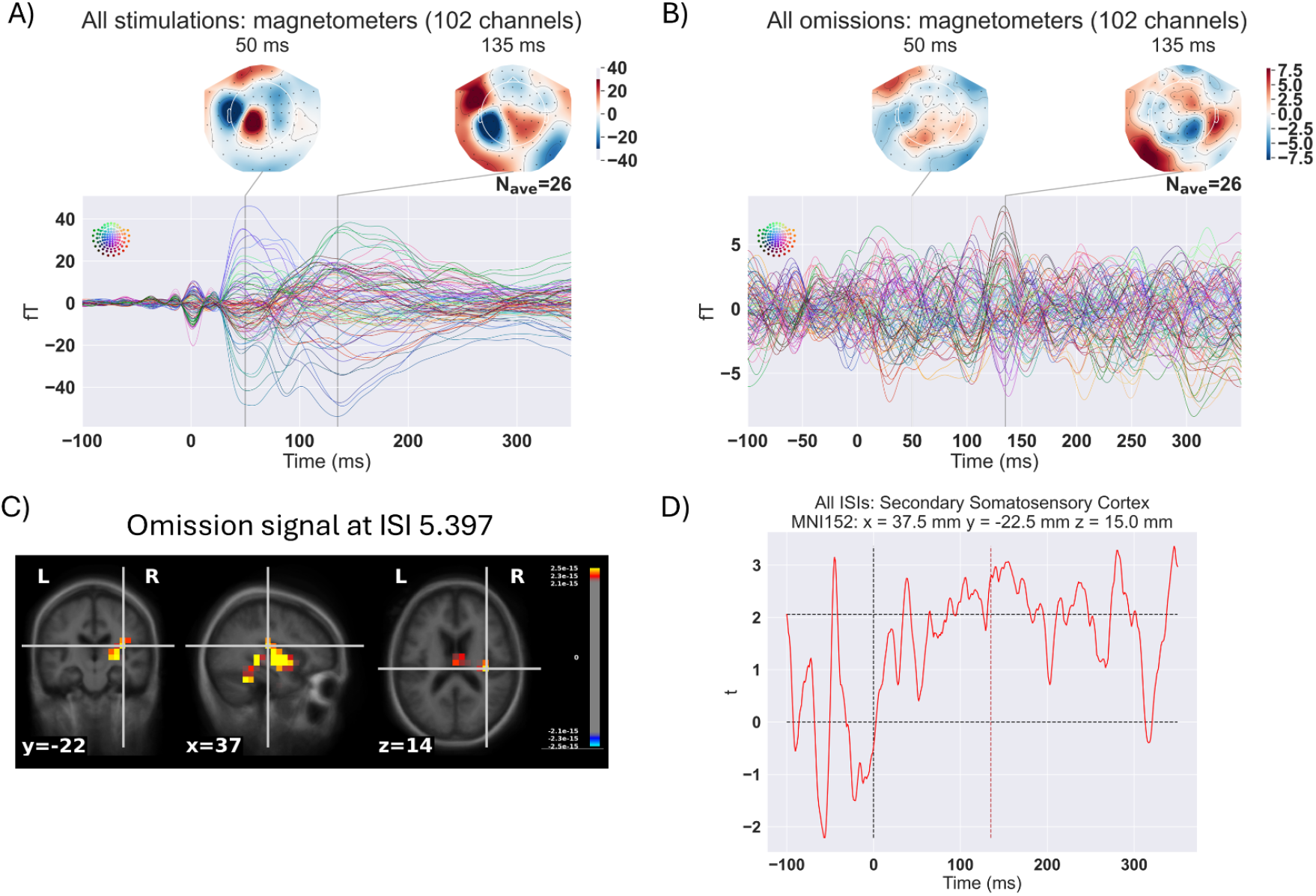
SII and SII results. A) The expected SI (50 ms) and SII responses (135 ms) are elicited for tactile stimulations. B) For omissions, no SI response is seen at 50 ms, whereas a SII response at 135 ms was detected. C) Source reconstruction of the omission signal (135 ms) for the longest ISI. The data shown are the 95th percentile for the whole brain at time 135 ms. The coordinate (*MNI152*: x = 37 mm, y = -22 mm, z = 14) is for the maximally responding source in the right secondary somatosensory cortex. D) The time course for the maximally responding source for all ISIs collapsed (at 135 ms, indicated with a red line) in the right secondary somatosensory cortex. The top black line indicates the *t-*threshold for significance with *α* = 0.05.

### 3.2 The Cerebellum

The analysis revealed a significant cerebellar response to omissions that was modulated by the temporal distance between stimulations. For the shortest ISI (0.497 s), tactile omissions elicited significant bilateral activation in cerebellar lobule VI, peaking at 33 ms post-stimulus at *MNI152*: *x* = 8 mm, *y* = -60 mm, *z* = -23 mm, *t*(25) = 3.35; *p* = 0.00257 (Figure 3A), indicating early cerebellar engagement in the detection of omissions. The cluster permutation test for this source showed a significant difference between the shortest ISI (0.497 s) and all ISIs collapsed, *p* = 0.039. The time range informing the rejection of the null hypothesis extended from 16 ms to 47 ms, part of the time range reported by Andersen and Dalal (2021). Having identified the peak value, we compared the shortest ISI to the remaining ISIs. For the contrasts between the shortest and the longest ISI (0.497 s vs 5.397 s), the same peak in lobule VI showed significantly increased cerebellar response to the shorter relative to the longer ISI in the time range from 16 ms to 47 ms (Figure 3B), *t*(25) = 3.37; *p* = 0.00244 (Figures 3B-C). The remaining tests showed: 0.497 s vs 4.597 s, *t* = 2.77; *p* = 0.0104; 0.497 s vs 3.397 s, *t* = 2.28; *p* = 0.0312; 0.497 s vs 2.597 s, *t* = 1.74; *p* = 0.0947; 0.497 s vs 1.397 s, *t* = 1.31; *p* = 0.204 (Supplementary Figure 2). This ISI-dependent modulation indicates that cerebellar lobule VI is more precisely engaged when stimuli arrive at shorter than at longer intervals, consistent with the cerebellum’s proposed role in generating precise temporal predictions.

**Figure 3.**
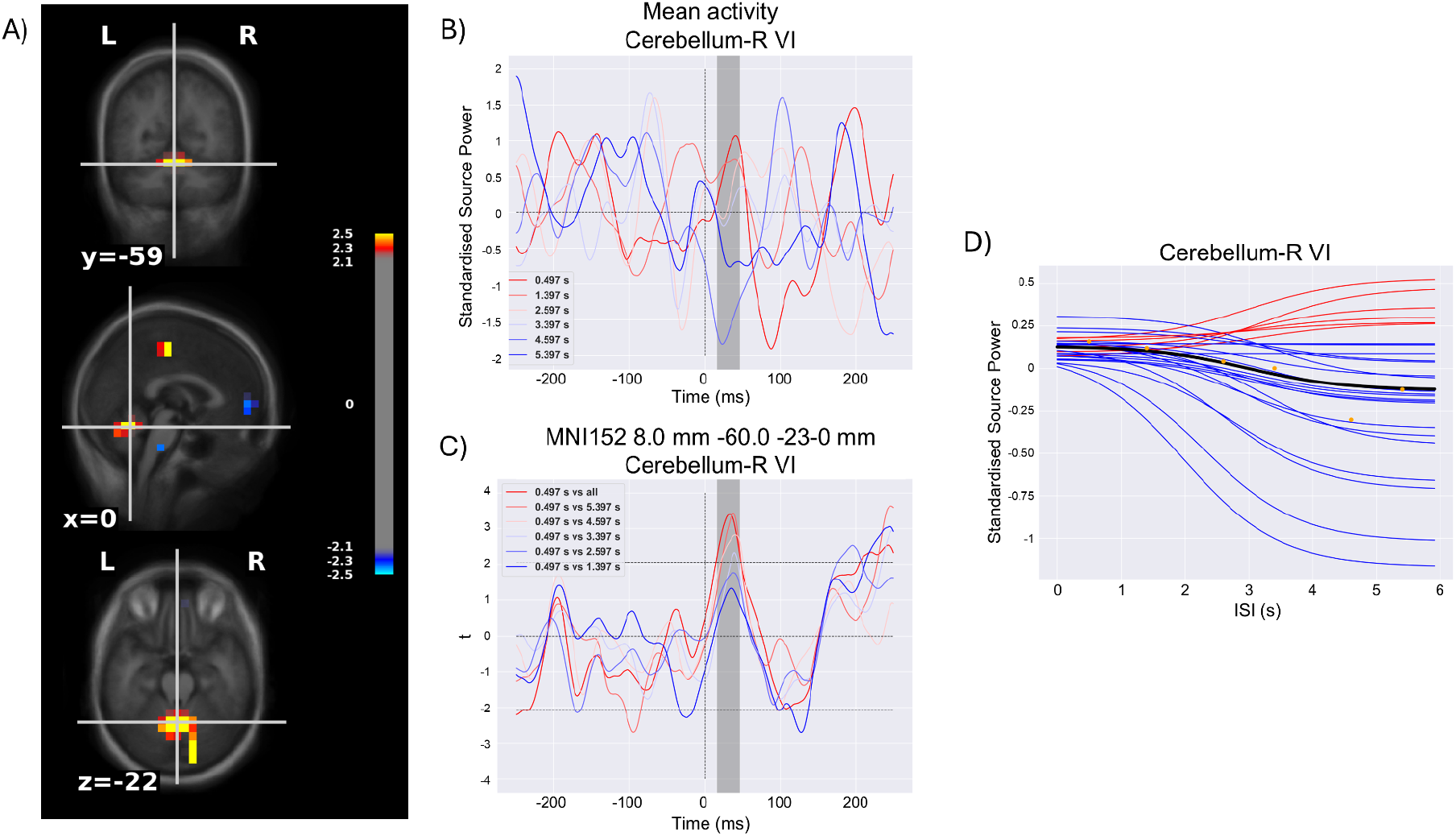
Cerebellar responses to omissions. A) Source space cerebellar results: statistical comparison between ISI = 0.497 s and all ISIs collapsed at 33 ms. B) Mean standardised activations of the peak cerebellar source, found in right cerebellar lobule VI, for the omissions. The stimulus should have been presented at t = 0 ms, if it had not been omitted. The grey box highlights the range, 16-47 ms, of the cerebellar response reported in panel A. Note how the 3-4 shortest ISIs cluster together on the positive side, whereas the 2-3 longest cluster on the negative side. C) *t*-values for comparing the shortest ISI (0.497 ms) against each of the remaining ISIs at the peak vertex. Note how *t*-values are higher when compared against longer ISIs in decreasing order. D) Fit of cerebellar response curve: the black line represents the mean of the posterior distributions for *a* (0.14), *b* (-0.04) and *c* (3.0 s) and a fixed *d* (0.75). Blue and red lines are fits for individual participants. Blue lines indicate participants whose *b* was less than their *a*, and vice versa for the red lines. Orange dots are mean values for each of the ISIs over the time range 16-47 ms.

To characterise the relationship between ISI and cerebellar engagement, we extracted mean neural activations from right cerebellar lobule VI for each ISI condition. The temporal profile revealed a step-wise decrease in cerebellar activation as ISI duration increased: shorter ISIs (< 3 s) elicited positive activation relative to baseline, longer ISIs (> 4 s) showed negative activation, while the intermediate ISI of 3.397 s exhibited activation levels between these two extremes. The estimation of the generalised logistic function (Equation 1) was compatible with a decrease in cerebellar activation to the omitted stimulus when ISIs became longer than 4 s, as was hypothesised (Figure 3D). The posterior distributions of the fitted parameters (Supplementary Figure 3) showed clear peaks for the upper asymptote (*a*), slope (*b*) and inflexion point (*c*) indicating reliable model convergence and constraint by the data.

## 4. Discussion

The results of this study underscore the importance of the cerebellum for temporal predictions within the somatosensory domain. Our findings reveal two key aspects of cerebellar engagement: the temporal precision of the prediction error response and the temporal boundaries of cerebellar predictive processing.

As a first step, we validated the omission paradigm by examining SI and SII responses. Neural activation to tactile stimulation was found in both SI and SII at the expected timepoints. For omissions, a clear SII response remained evident with a topographical distribution consistent with contralateral SII activation (Andersen & Lundqvist, 2019).

When collapsing across all ISIs, we observed a cerebellar response to omissions peaking at 30–40 ms after the expected stimulus, consistent with previous findings (Andersen & Dalal, 2021, 2024). Crucially, we found that the magnitude of cerebellar activation during stimulus presentation was strongly modulated by temporal predictability, with lobule VI showing greater engagement for shorter ISIs compared to longer ISIs, thus corroborating our hypothesis.

The consistency of the timing of this response in the current study and previous work suggest that the cerebellum is highly attuned to deviations from expected timing, likely reflecting its function in generating and maintaining internal models of temporal regularity. Furthermore, the magnitude of the cerebellar omission response was compatible with a logistic decay pattern, diminishing for ISIs exceeding 4 seconds (Figure 3D). The inflexion point of this decay curve (*c*) fell within the 2–4 second window, consistent with the framework proposed by Tesche & Karhu (2000), indicating a decline in the cerebellum’s predictive accuracy beyond this temporal range. It must be noted that the prior on *c* was not changed considerably by the data, meaning that the posterior was very similar to the prior. While the group-level logistic decay curve shows a clear inflexion point around 2–4 seconds, individual participant fits (Figure 3D) reveal heterogeneity in asymptote values (*a & b*). 6 participants exhibited flat or inverted patterns, showing greater cerebellar activation at longer ISIs than at shorter ones. The group-level effect, however, is driven by the majority of the participants (n = 20) who show the hypothesised pattern. The origins of this variability, whether reflecting genuine individual differences in cerebellar temporal processing capacity or methodological constraints, remain an open question.

While the main hypothesis was corroborated, the precision with which this boundary can be determined remains an important limitation of this study. The experimental design employed equal trial sampling across all six ISIs, rather than denser sampling around the expected inflexion point. Consequently, the data provide limited constraint on the precise location of this parameter, rendering the estimate sensitive to the prior specification in the model. The 2–4 second window should therefore be interpreted as a broad approximation rather than a precise temporal threshold. Additionally, fixing the steepness parameter *d* constrains the model to a step-like transition, which may not accurately reflect the underlying neural dynamics. As with the inflexion point, the sparse sampling around the 2–4 second window provided insufficient constraint to reliably estimate the steepness of the transition, preventing *d* from being estimated freely. With richer sampling around the inflexion point, future research could empirically test whether a logistic decay pattern is the best approximation of the neural response. A natural progression of this work would be to include behavioural measures to examine whether these time-constrained cerebellar predictions translate into measurable effects on perception. For instance, investigating whether detection sensitivity or reaction times to unexpected stimuli scale with ISI in a manner consistent with the logistic pattern observed in the neural data.

In conclusion, this study provides evidence that the cerebellum plays a central role in temporal prediction within the somatosensory domain, operating within specific temporal constraints. The robust cerebellar responses to omitted stimuli at the precise latency when stimulation was expected, combined with the systematic modulation of cerebellar activation by ISI duration, demonstrate that lobule VI maintains temporally precise internal models of expected sensory timing. The logistic decay of prediction responses beyond 2–4 seconds is compatible with an empirical boundary for cerebellar predictive processing. However, the precise location and sharpness of this transition remain to be established in future work. Together, these findings support a model in which the cerebellum generates precise temporal predictions that are transmitted to higher-order sensory areas such as SII, enabling the brain to anticipate the timing of upcoming events and rapidly detect deviations from expected temporal patterns. This predictability sensitivity is consistent with the broader literature indicating that the cerebellum is crucial for integrating past regularities to form predictions about when sensory events should occur, and for signalling when those predictions are violated (Breska & Ivry, 2021).

## 5. Data and Code Availability

The code that supports the findings of this study is openly available on GitHub https://github.com/ualsbombe/temporal_brains. Data cannot be shared, as we cannot guarantee the anonymity of participants if magnetic resonance images are shared. MEG data can be shared upon reasonable request.

## 6. Author Contributions

K.H.: Software, Methodology, Formal analysis, Investigation, Data curation, Writing – original draft, Writing – review & editing, Visualisation, Funding acquisition. L.M.A.: Conceptualisation, Methodology, Software, Validation, Formal analysis, Investigation, Resources, Data curation, Writing – original draft, Writing – review & editing, Visualisation, Supervision, Project administration, Funding acquisition.

## 9. Acknowledgements

The authors wish to thank S.W. Nehrer and M.M. Hansen for their valuable assistance with data collection.

## 11. Supplementary Material

**Supplementary Figure 1.**
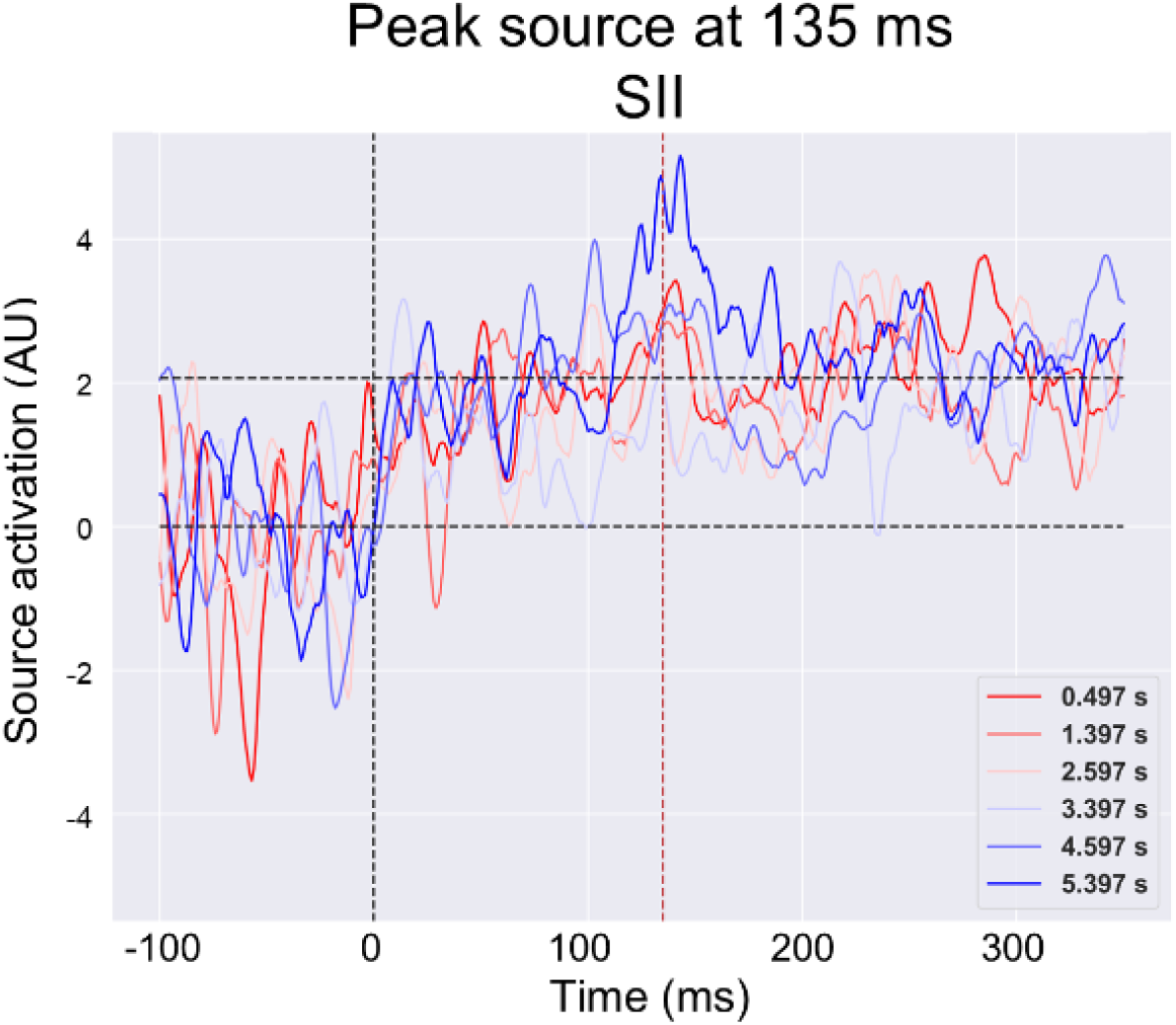
SII responses to omissions. The maximally responding sources (at 135 ms) in SII to the different ISIs. Notably, the strongest response was to omissions of the longest ISI (5.397 s). The top black line indicates the *t-*threshold for significance with *α* = 0.05

**Supplementary Figure 2.**
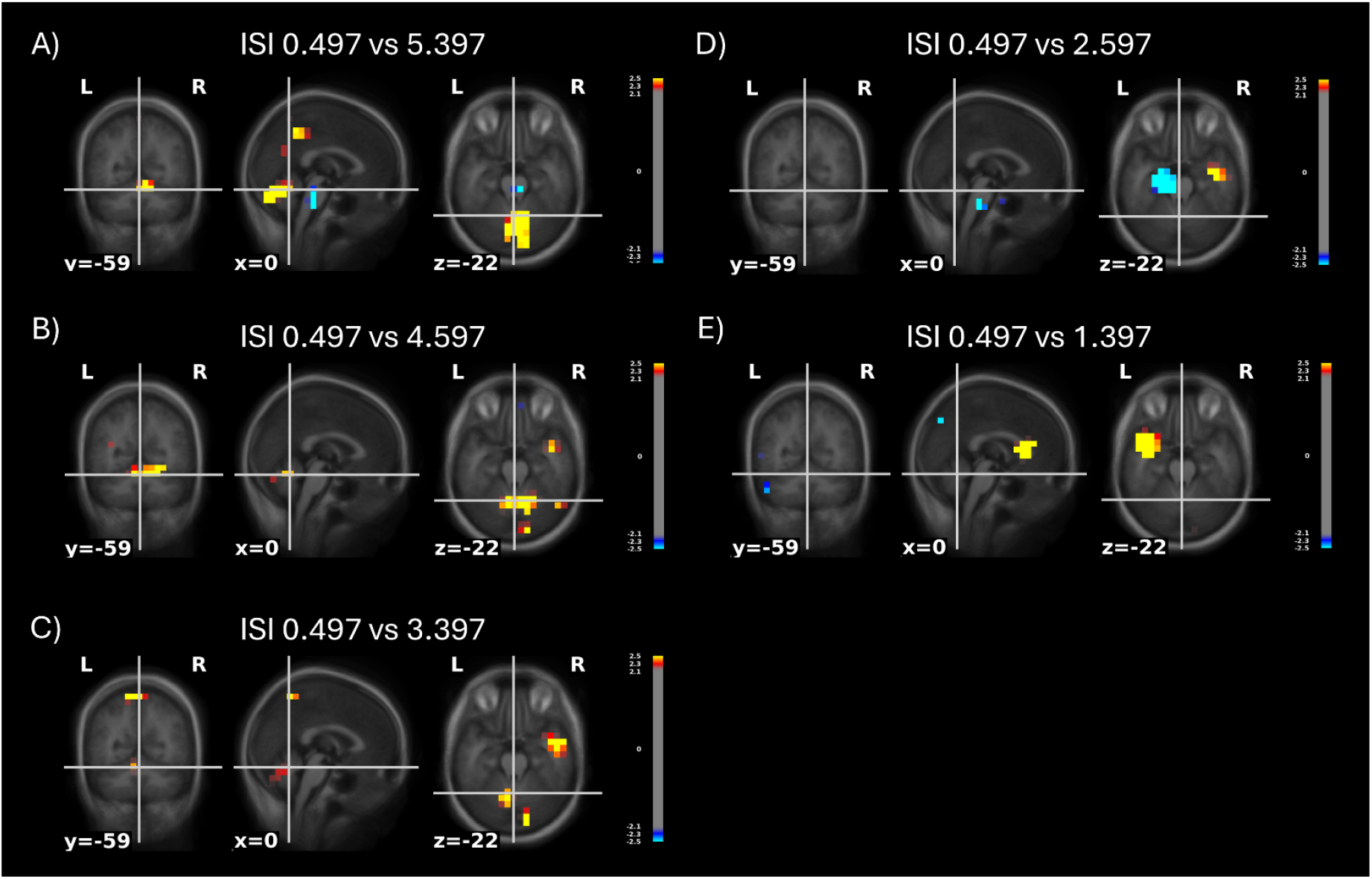
Source space comparisons of the shortest ISI to all other ISIs individually. Each panel shows a comparison between the shortest ISI (0.497 s) and the remaining ISIs (5.397 s, 4.597 s, 3.397 s, 2.597 s, 1.397 s, in that order).

**Supplementary Figure 3.**
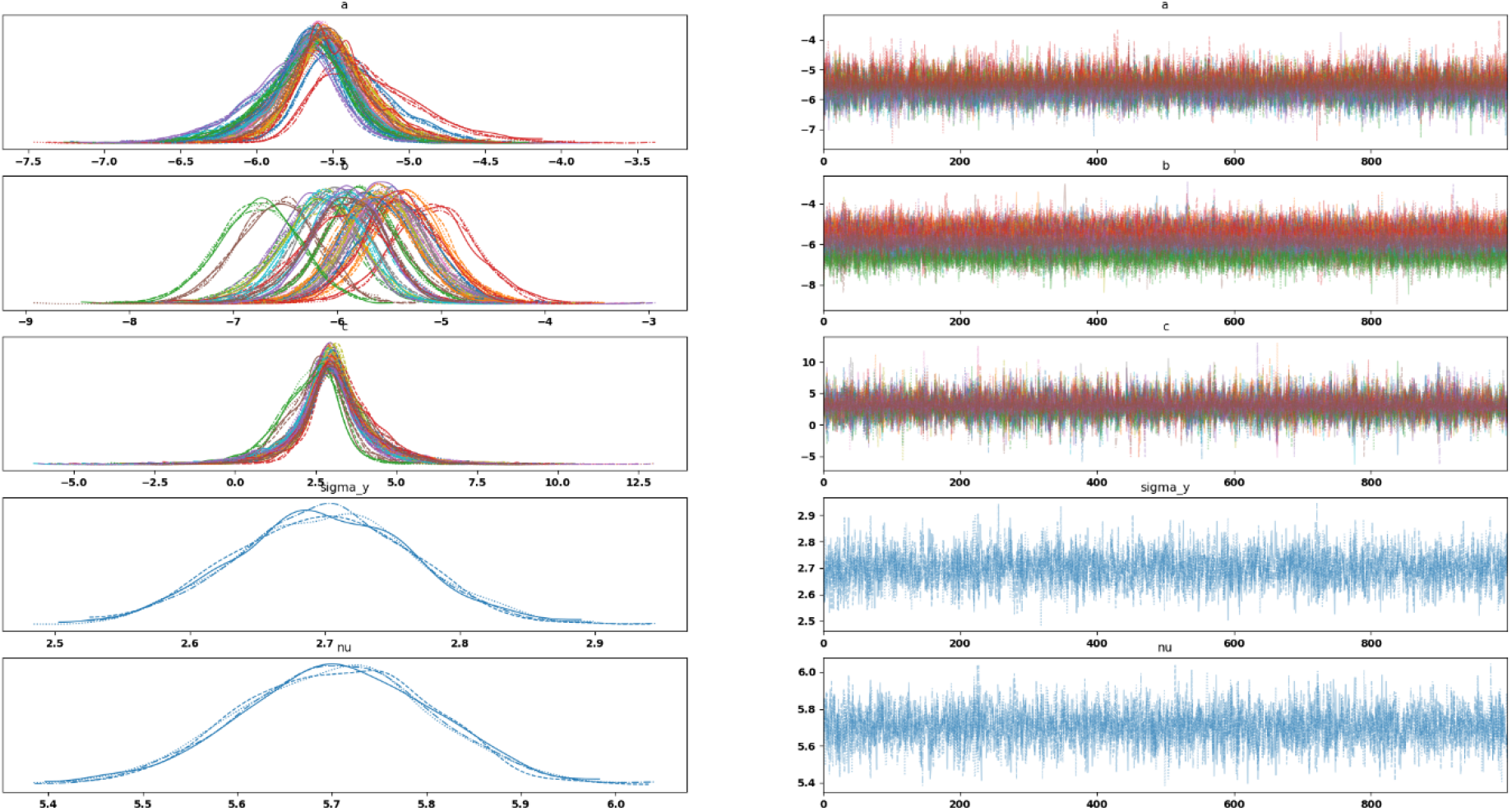
Posterior distributions of fitted parameters for the cerebellar response curve. The figure shows posterior distributions for each parameter of the generalised logistic function fitted to cerebellar omission responses: upper asymptote (*a*, mean = 0.14), slope (*b*, mean = -0.01), inflexion point (*c*, mean = 3.0 s), and the steepness parameter (*d*, fixed at 0.75).

